# Perception is Rich and Probabilistic

**DOI:** 10.1101/2022.07.03.498587

**Authors:** Syaheed B. Jabar, Daryl Fougnie

## Abstract

When we see a stimulus, e.g. a star-shaped object, our intuition is that we should perceive a single, coherent percept (even if it is inaccurate). But the neural processes that support perception are complex and probabilistic. Simple lines cause orientation-selective neurons across a population to fire in a probabilistic-like manner. Does probabilistic neural firing lead to non-probabilistic perception, or are the representations behind perception richer and more complex than intuition would suggest? To test this, we briefly presented a complex shape and had participants report the correct shape from a set of options. Rather than reporting a single value, we used a paradigm designed to encourage to directly report a representation over shape space—participants placed a series of Gaussian bets. We found that participants could report more than point-estimates of shape. The spread of responses was correlated with accuracy, suggesting that participants can convey a notion of relative imprecision. Critically, as participants placed more bets, the mean of responses show increased precision. The later bets were systematically biased towards the target rather than haphazardly placed around bet 1. These findings strongly indicate that participants were aware of more than just a point-estimate; Perceptual representations are rich and likely probabilistic.

## Introduction

Imagine being shown a star-shaped object. After taking it away, what awareness of it would you have? Would you be aware of having seen its shape, or would your awareness of it be hazy and probabilistic? Our intuitive idea is that perception would be discrete^[1]^: Either we would see the star, or if the percept was too brief, perhaps nothing at all. However, the machinery of perception produces probabilistic computations and mechanisms throughout the visual system. For example, orientation-selective neurons in primary visual cortex have a preferred orientation with the firing rate of neurons dropping off the further the stimulus is from their preferred orientation^[2]^. The same also applies to moving stimuli in the MT cortex^[3]^ and neural firing in response to visual stimuli in general resembles probability density functions^[4]^. Decoding of these neural activities, both in visual and prefrontal cortices, demonstrate correlations with reported confidence^[5]^, suggesting that people are aware of and can report their internal representations at least to some extent. Given that neural representations are probabilistic, how could perception be discrete? Indeed, visual perception may only seem non-probabilistic since our vision is rarely imprecise. If the star shape is seen just for an instant, maybe then we might feel unsure if asked to report on the number or depth of the points. Such a feeling could reflect that the underlying representation is hazy or probabilistic.

Our contention is that perception is rich, complex, and probabilistic. It is “rich” in that representations are more than a discrete point^[1]^, and “complex” in that it is more than just discrete representation coupled with some sense of confidence (e.g., a symmetrical interval or Gaussian around the discrete point). “Probabilistic” we take to mean that the internal representation is akin to a distribution over a feature space (a shape space, in this case).

Although the computations that assist vision are probabilistic^[6,7,8,9,10,11]^, several researchers have argued that the outcome of perceptual computations, while probabilistic themselves, result in single, discrete percepts. For example, Block^[1]^ proposes that “competition among unconscious representations yields conscious representations through winner-takes-all processes of elimination or merging”. In support of such a view, researchers have found that perceptual decisions show little evidence of knowledge of the probability distributions formed during perception^[12]^. Other studies reveal conflicting results, but since they use an array of target stimuli per display, e.g. orientations that have some spread^[13,14]^, they do not meet the ideal criteria for isolating perception. For example, Rahnev and Block^[15]^ argue that perception is only likely to be discrete in instances in which an object is briefly presented in isolation. Otherwise, probabilistic computation could arise from multiple discrete perceptual events being combined at decision-making stages.

The advantage of perceptual decision-making tasks is that they reveal what properties of a perceptual representation are available to higher-level cognition. However, a disadvantage is that if decisions are not using encoded information optimally it is unclear whether perception lacks this information or whether this information is not incorporated into decisions. Therefore, our task was designed to allow participants to directly report their internal representations rather than inferring representational properties from the optimality (or sub-optimality) of decisions. We used a modified continuous report paradigm—where participants report the value of a shown shape from a continuous shape space arrayed on a circle—to allow participants to place ‘bets’ over the circular feature space. Participants placed multiple Gaussian bets over positions on the circle and the combination of these bets allowed participants to draw an ‘uncertainty profile’ on a per trial basis.

To preview our results, we found evidence of rich and probabilistic information in the reports. The spread of the uncertainty profiles, which can be taken as a measure of representational noise, was correlated with the imprecision of reports, indicating an awareness of the amount of imprecision in their representations. The drawn uncertainty profiles matched the shape of bet 1 errors, further demonstrating that later bets within a trial were likely constrained by representational content rather than reflecting strategy. Most importantly, while the later bets in a trial were associated with larger error magnitudes than bet 1, they cumulatively shift the mean of the uncertainty profile closer and closer to the true stimulus. The later bets were systematically biased towards the target rather than haphazardly placed around bet 1. Taken together, these findings point towards participants having an awareness of the target that extends beyond one discrete response or even a discrete response plus a separate notion of confidence^[15]^.

## Methods

### Participants

Forty naïve participants (16 female, median age 22) were recruited for Experiment 1. Our sample size was based on previous work—a previous study on working memory^[16]^ using a similar paradigm had sufficient power with 40 participants to detect the effects of interest. All participants declared normal or corrected-to-normal visual acuity and color vision. Participants were recruited online (via www.prolific.co) for a base allowance of 5.20GBP per hour and were told that they could also receive a monetary bonus depending on their performance (mean bonus = 2.46 GBP). The experiments were approved by the New York University Abu Dhabi Institutional Review Board and carried out in accordance with relevant guidelines and regulations. Informed consent was obtained from all participants.

### Apparatus and stimuli

Because of the online nature of the experiment, stimulus sizes would vary slightly depending on the participant’s screen size/viewing distance. Assuming that a 27-inch 1920 × 1080 resolution monitor placed 70 cm away was used, targets would be from the center of the screen, with the target shapes themselves occupying 2° of visual angle. The background of the screen was white throughout the experiment. Targets (one per trial) always appeared off-center from the fixation point (approximately 2° away), at some random bearing. Effort was made to ensure that the drawn display fit within the participant’s browser page. Participants were also asked to maximize their browser window prior to launching the experiment. All reported experiments were programmed entirely in HTML Canvas/Javascript.

The targets were derived from the validated circular shape space^[17]^, which consists of 360 shapes (one per integer degree angle of the circle). As with typically used color wheels, each shape was only slightly different from its neighbors and angular distance along its 2D circle is correlated to visual similarity (Figure 1a), resulting in a continuous and circular stimulus space. Stimulus display duration was 40ms. The targets were degraded/noised by replacing 90% of the black pixels with white ones to match the background (Figure 1c). To recap, from our original design to investigate working memory^[16]^ we made three important changes overall to adapt it to study perception, first was the switch to a single shape stimulus per display, not an array^[13,14]^, since having multiple items introduces problems with confusion among the items^[18,19,20,21]^. Second was the short stimulus timing, and third was the noise added to the stimuli. These latter two were to increase the likelihood that we still have performance below ceiling while having only one stimuli per display.

**Figure 1.**
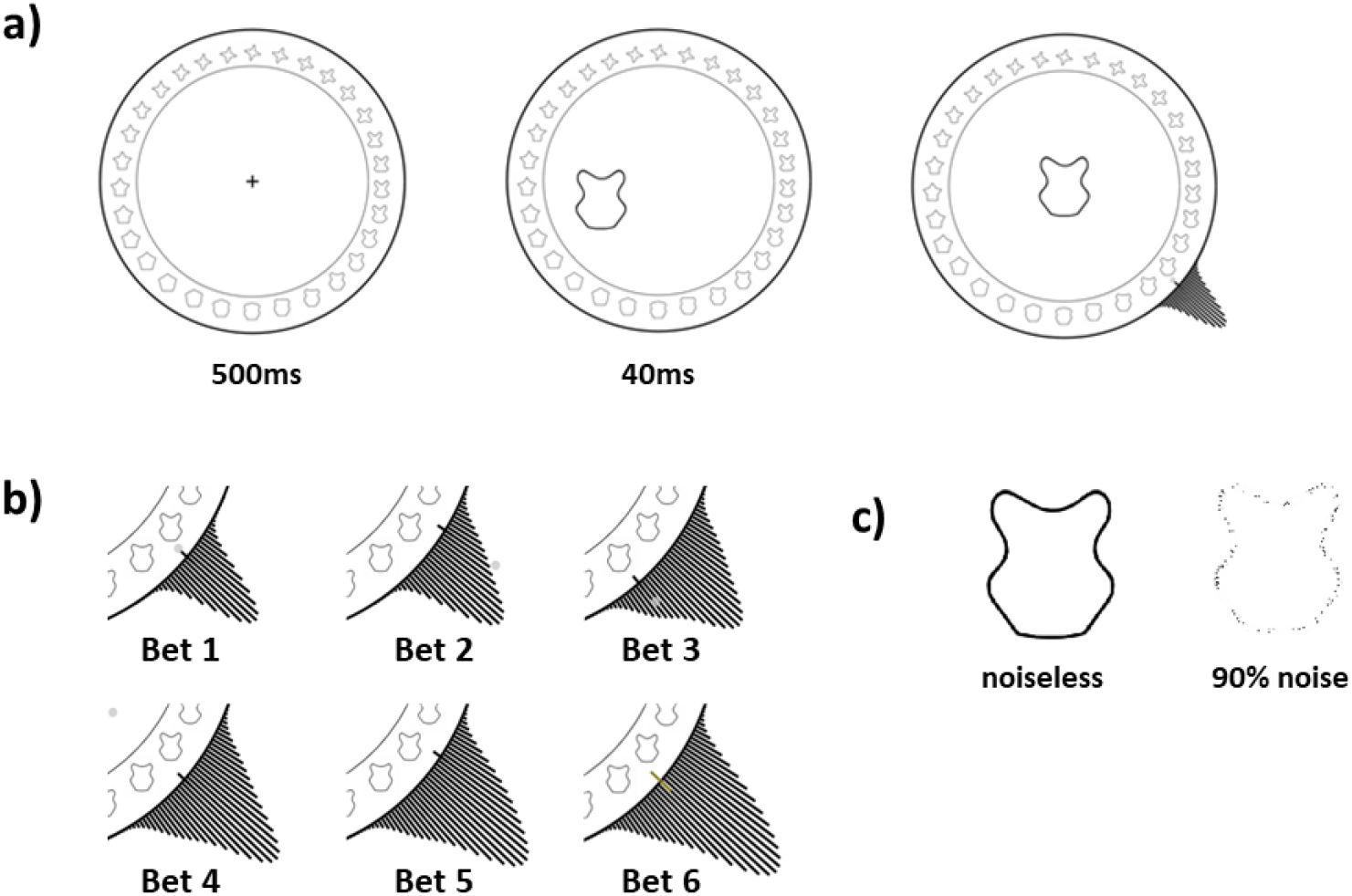
Experiment Paradigm. a) Trials began with a 500ms fixation. A target shape then appeared (at some random bearing off-center) for a short duration of 40ms. For illustration the target is noiseless, but actually was noised in the experiment. Thereafter, the participant is free to make their bets on what they saw using the shape wheel. The currently selected shape appeared on the center of the screen. b) Participant bets add Gaussian (*SD* = 4°) component on the circle, which was always previewed to participants pre-click. Participants were awarded points based on the height on the target shape’s position on the circle. Participants were given incentive to spread their bets as stacking provided diminishing returns on height. c) Illustration of the noise effect of replacing the black pixels with the background color. The shapes and shape wheel were adapted from Li et al.^[17]^.

### Procedure

Each trial began with a blank screen with a fixation cross lasting for 500ms. A randomly selected noised target shape then appeared on-screen for 40ms, at some off-centered bearing from fixation. Stimulus presentation was *immediately* followed by the response phase.

Participants were asked to indicate the shape that they remembered using the shape wheel (see Figure 1a). The chosen shape was previewed to participants and updated at the center of the wheel with every mouse movement. A black Gaussian distribution (standard deviation of 4° on the shape space) also appeared on the wheel, centered on and updated with participants’ mouse location. Participants confirmed their choice with a mouse click.

For the 2nd to 6th bets, a similar Gaussian appeared at a random position on the bar, and participants were asked to move the mouse to place the Gaussian over the shape they want to respond with, again confirming the bet with a mouse click (iteratively, for a total of six bets). Note that the Gaussian for bets 2-6 had half the height of the Gaussian for bet 1. This was to encourage participants to be as accurate as possibly during the first bet. During each bet, the uncertainty profile (based on current mouse position) was updated per frame and previewed to participants. Participants were instructed to build a distribution that matched their internal state. The six Gaussians summed to create a final uncertainty profile for that trial. Participants were free to stack responses on top of one another or to spread it across a larger shape space. As a result, the uncertainty profile ‘drawn’ at the end of the sixth response need not have a Gaussian shape. Note that the Gaussians spawned in a random location between responses to prevent stereotyped successive clicks on the same shape location, and participants had to wait a minimum of 300ms and make a lateral mouse movement before they could register the next click. To encourage participants to report something resembling the true internal representation, participants were awarded points based on the height of the final drawn distribution over the target shape:

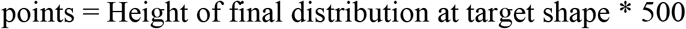

The first bet was twice as tall (i.e., worth twice as many points) to encourage participants to place the first bet accurately. To prevent participants from adopting the strategy of stacking responses at the first bet, we implemented diminishing returns to stacking by reducing the value or rewarded points when stacking bets on existing bets. This penalty was scaled by height. Specifically, the height of the Gaussian component to be added was penalized by the current height of the uncertainty profile raised to a penalty parameter. For example, if the current bet is b, and the uncertainty profile already drawn due to the previous bets is y, then the new bet y’ is:

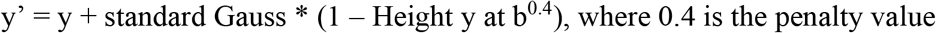

The taller the current height at the new bet value, the smaller the possible gain in height (and therefore the smaller the gain in points). Importantly, this penalty was built into the visualization of the drawn distribution seen by participants. After the six guesses were placed, the trial score, total score, as well as the current accumulated monetary bonus (every 200 points earned 0.10 GBP) were displayed on the feedback screen until the participant re-centered the mouse on the central fixation. The maximum possible points per trial was 200. Each participant first completed 6 practice trials, followed by 150 main trials, with a break every 50 trials. Practice trials were excluded from the analyses.

## Results

### Evidence that the reported distributions incorporate imprecision

To explore the degree to which participants’ bets captured representational noise, we first looked at whether the spread of bets predicted performance. To maximize performance, a participant should place bets narrowly when the shape representation is precise is low and spread bets widely when precision is low. Indeed, we found significant positive correlations (*p*< .05) between the magnitude of the error of the first response and the median absolute distance between adjacent bets for 33 of the 40 participants (mean *r* = .392). These individual correlations were Fisher-transformed (mean *z* = .434, *95% CI* = [0.029, 0.899]) and a *t*-test found this distribution to be significantly different from zero (*t*(39) = 11.75, *p* <.001). Similar results were found using other measures of the bet spread, including the standard deviation (mean *z* = .343) or the interquartile range^[22]^ (mean *z* = .333) of the uncertainty profile.

### Evidence that profiles reflect trial-specific probability

This analysis demonstrates that the way the bets are placed reflects the error in participants’ first response. But how closely does the reported uncertainty profiles within trials match participants’ across-trial error, as assessed by the uncertainty profile of the error in first responses across trials? If participants were accurately recreating their internal representations by sampling from it to derive first responses, the across trial error distributions would match the average of the uncertainty profiles reported within a single trial. We examined this using two-sample Kolmogorov-Smirnov (*KS*) tests^[22]^ to compare the across-trial error distribution (error of the first response relative to the correct answer) to the trial averaged uncertainty profile for each participant. Uncertainty profiles were circle-shifted to align the target shapes and averaged at each integer value of the shape space (e.g., at 360 discrete points). In 32 of the 40 participants, the *D*-value (the *KS* statistic which measures the maximum difference between the two empirical cumulative distribution functions) was small (mean *D* = .045, *SD* = .025) and non-significant (*p*>.05), highlighting that the average reported uncertainty profiles was not significantly different from the across-trial error of the first response (Figure 2a). This analysis suggests that participants have access to, and can report, rich probabilistic information. The manner in which participants spread their bets are likely constrained by their internal representations.

**Figure 2.**
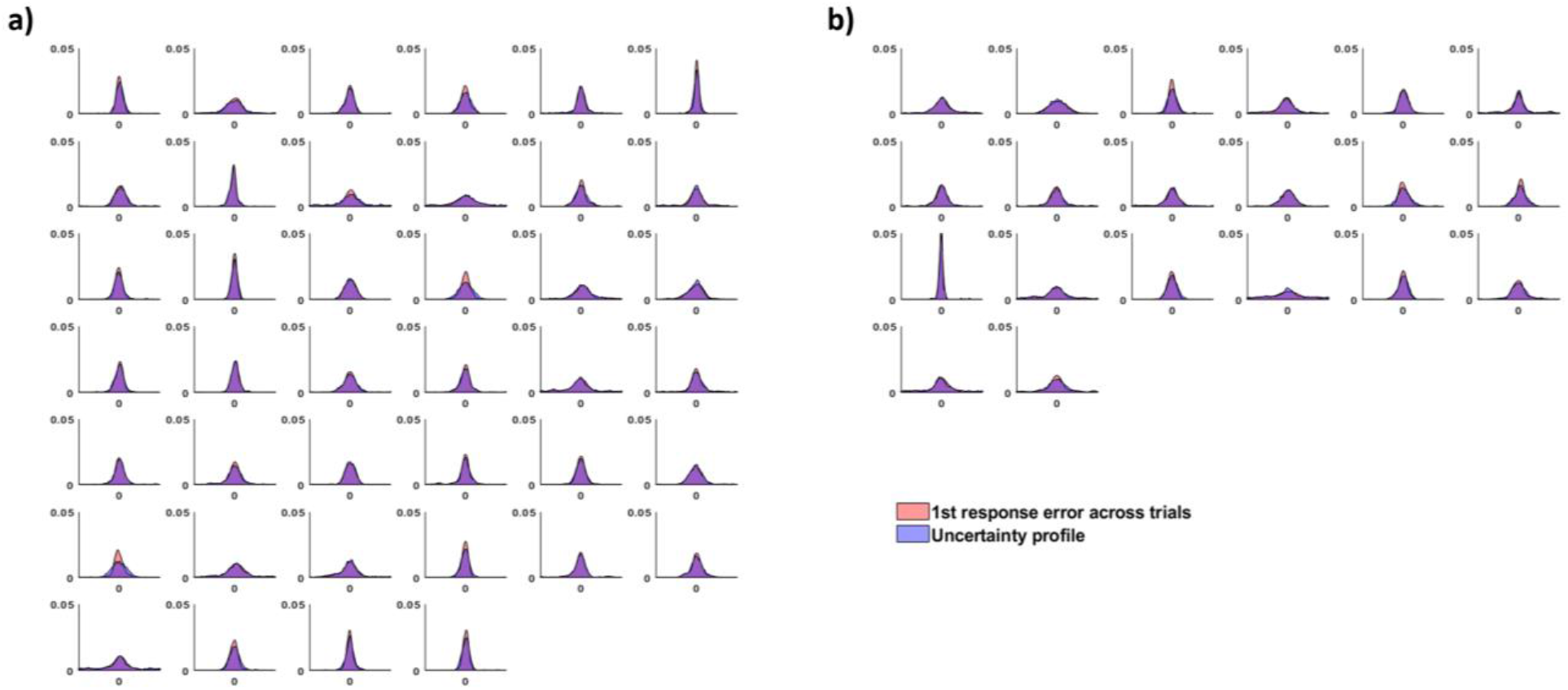
Comparing the error distribution of the first response across trials (red) versus the drawn uncertainty profiles (after all six bets) averaged across trials (blue), for each participant. a) The 40 participants of in the main study. b) 20 participants from a control study where participants had the ability to undo a response.

Further, if participants were just using a confidence plus a discrete remembered percept^[15]^ rather than having access to a rich internal distribution, we would expect that the drawn distributions should be roughly symmetrical about bet 1. Therefore, if we flip the uncertainty profiles before comparing it against the across-trial error distribution, it should not significantly affect the *KS* test results. However, doing this results in only 11 out of the participants (versus 32 in the original analysis) having the two distributions be non-significantly different (mean *D* = .098, *SD* = .064). Comparing this to the original analysis, these *D*-values are significantly different (*t*(39)=5.05, *p*<.001). In other words, the overlap between the averaged within-trial uncertainty profiles and the across-trial first bet error density plot is significantly worse if we flip one of the two distributions, suggesting that there is important information about the target percept held in the asymmetry of the uncertainty profiles.

### Evidence that the bets contain more information than found in the first response

The previous analyses highlight that the uncertainty profiles convey rich information about internal representations. But do they also contain information about what is in memory beyond what is captured by the initial response or the first response plus a separate notion of confidence^[15]^? If participants have an internal distribution from which to make multiple responses from, it is possible that the cumulative error across bets is more precise than any single bet, even the first one (see Supplementary Materials). An example of how this could work is if participants have an internal representation but are able to only draw one sample from it at a time (e.g., one different sample per bet), rendering the initial bet suboptimal. This is unlike a discrete or a discrete + confidence model, where the cumulative error can only get worse (particularly if we assume there is some memory component that is causing a decay of information over the bets).

We analyzed the placement of individual bets, as well as the cumulative circular average of the bets (e.g., if the first bet was a shape 10° clockwise and the second was 4° counterclockwise, the cumulative average would be 3° clockwise). As shown in Figure 3, the first bet placement is the most accurate, with a relatively monotonic decline in the accuracy of individual bets across the trial (when considering the center of bets in isolation). To examine this, we subjected the bets to a one-way *ANOVA* with the six levels of bet order as a within-subjects factor. Errors generally increased with bet number (bet 1 = 14.9°, bet 6 = 20.3°), *F*(5,195) = 17.76, *p* <.001, *η*^2^ =.313. This result is not surprising given that the first response was worth double the points of subsequent responses and that subsequent responses occurred after increasing delays and potential interference from previous responses.

**Figure 3.**
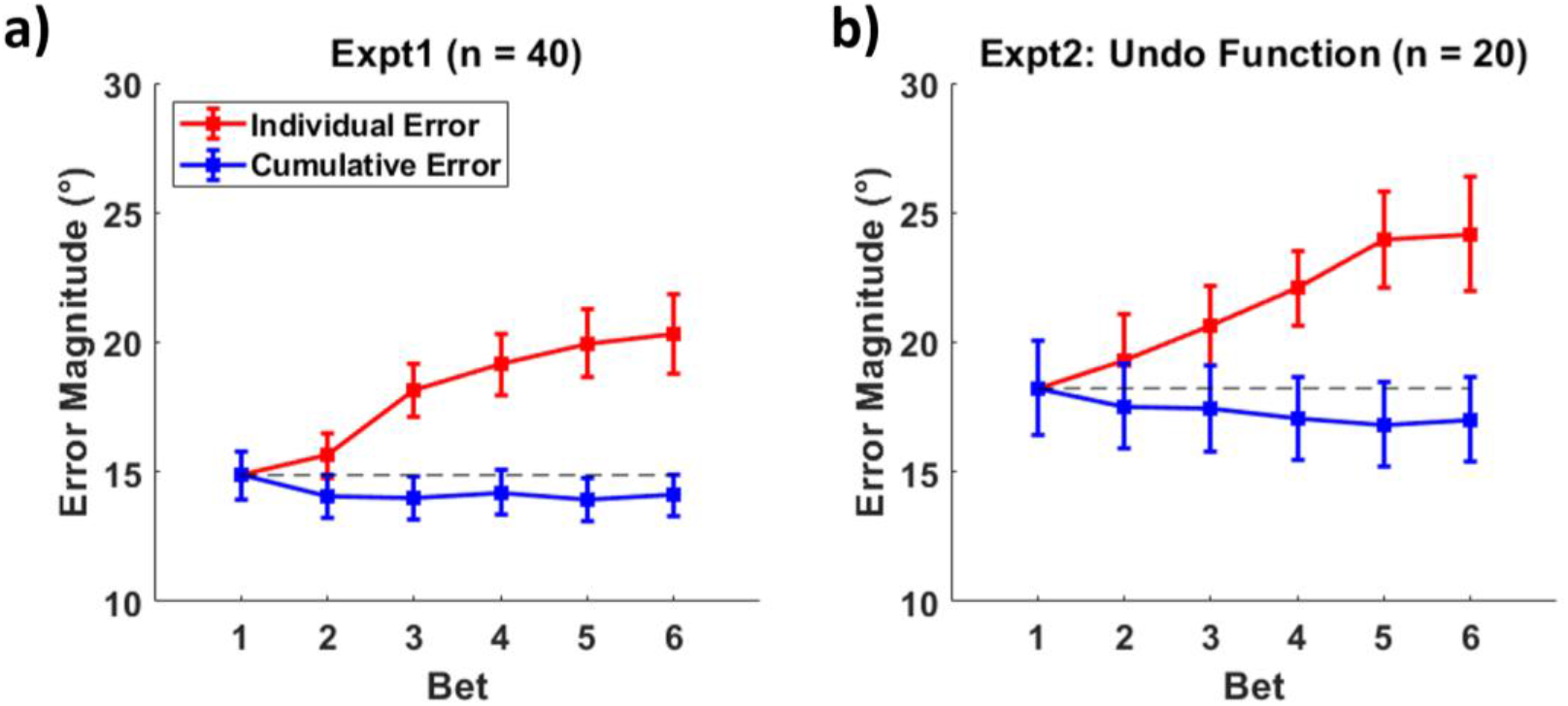
Individual response errors (red) and cumulative errors (blue) as a function of response order. Cumulative errors are calculated as the error of the mean of responses. For example, the cumulative error for response 3 would be influenced by response 2 and response 1, whereas the individual response error for response 3 is solely determined by that bet. a) Errors for Experiment 1. b) The equivalent errors for Experiment 2.

Despite being less accurate than the first responses, subsequent responses could nevertheless contain additional, unique information about the target item. To test this, we examined whether additional bets were providing novel information not contained in the first response by testing whether the center of the uncertainty profile approaches the true shape with repeated placement of bets. We subjected mean bet placement to a one-way *ANOVA* and we found that performance improved with additional bets (bet 6 = 14.1°), *F*(5,195) = 3.71, *p* = .003, *η*^2^ =.087, despite the fact that average error trended much worse for the later bets. This suggests that the combination of responses contains more information than any individual response, including the first response.

### Evidence that averaging benefits are not caused by errant first responses

As a precaution, we wanted to ensure that any effects were not due to participants making a mistake at Bet 1 (e.g., a mis-click) but with correct reports in subsequent bets^1^. The paradigm for this control study was virtually identical to Experiment 1, except that for any given trial, participants were given the ability to erase one bet by making a right-mouse click. For example, if they made a mistake for Bet 1, then can go back before confirming their Bet 2. Likewise, a participant can undo Bet 5 before confirming Bet 6. Participants who did not use this feature within the first 20 trials were given a reminder of this functionality. All participants were also reminded of it between the practice/main trials and on each break.

Participants rarely used the undo function (mean = 2% of trials), despite the constant reminders. Pointing-related error rates show a speed-accuracy trade-off^[24]^; thus, a low-correction rate was to be expected given the non-speeded nature of the task. These undone bets were generally of greater error magnitude error (mean = 70°) and in-line with random clicking (chance error = 90°) compared to the bets that replaced them (mean = 16°), indicating that these were mis-clicks rather than true responses. The use of the undo function was split evenly across the bets, e.g., it was used as frequently for undoing bet 1 as it was used for bets 2, 3, 4 or 5. Still, an independent *t*-test on the Bet 1 error magnitudes for Experiment 2 (*M* = 18.2°, *SD* = 1.3°) was marginally higher than the one for Experiment 1 (*M* = 14.9°, *SD* = 0.9°), (*t*(58) = 1.82, *p* = .074, *η*^2^ = .05) despite the ability to erase an incorrect response, suggesting that Experiment 1 did not suffer from much of these mis-clicks.

Otherwise, results of this control study mirror that of the main study. Significant positive correlations (*p*< .05) were also found between the magnitude of the error of the first response and the median absolute distance between adjacent bets for 19 of the 20 participants (mean *r* = .440). When comparing the error distribution of the first response across trials versus the uncertainty profiles (after all six bets) averaged across trials (Figure 2b), in 14 of the 20 participants, the *KS* test *D*-value was small (mean *D* = .055, *SD* = .016) and non-significant (*p*>.05). Errors also generally increased with bet number (bet 1 = 18.2°, bet 6 = 24.2°), *F*(5,95) = 11.82, *p* <.001, *η*^2^ =.384, while the cumulative bet error (bet 6 = 17.0°) decreased, *F*(5,95) = 2.51, *p* = .035, *η*^2^ =.12 (Figure 3b). Therefore, we see no evidence that the improvement from later responses is caused by participants realizing after they have made an errant response.

## Discussion

We rapidly perform complex calculations and process information about our environment. The neural computations supporting perception are capable of producing probabilistic information, but is this information available to consciousness and higher-level decision-making? This has generally been studied by determining whether humans can perform flexible and deliberate computations that require access to more than summary statistics^[15]^. Studies using this approach have yielded mixed results^[12,13,14]^ and there are concerns about whether a failure to perform optimal calculations reflects a lack of information about internal representation, or just an ability to effectively reason about it. Here we try a more direct approach by asking participants to place multiple Gaussian bets per trial over a shape space in order to convey a sense of probability of a perceived shape. The summation of these Gaussian bets create uncertainty profiles on a per trial basis. Several lines of evidence suggest that participants have access to more than a discrete shape and can report something akin to probability distribution. Spread of the bets/ uncertainty profiles was correlated with first bet precision, indicating an awareness of the amount of imprecision in representations. The uncertainty profiles matched the shape of bet 1 errors, further demonstrating that later bets within a trial were likely constrained by internal representations rather than reflecting strategy. Most importantly, while the later bets in a trial were associated with larger error magnitudes than bet 1, they cumulatively shift the mean of the uncertainty profile closer and closer to the true stimulus. This is unlikely the case if participants were only using some sense of confidence to guide how to spread the later bets: Because bets would be spread symmetrically about bet 1, cumulative error should not decrease). All these point towards participants having an awareness of the target that extends beyond one discrete response. Using the betting game paradigm, we similarly found evidence that working memory representations were rich^[16]^.

It may not seem surprising that the betting game task would yield similar results when applied to perception or working memory, given that there is considerable evidence that working memory representations are perceptual representations that are maintained by internal attention^[25,26,27]^. However, this is not to say that the two tasks would be driven by the same representations. Indeed, there were major differences in the two tasks: stimulus presentation (single-item versus an array), task demands (retention interval or instant report) and performance (reports for working memory tasks have considerably more variability). Rather, this suggests that the representations that occupy the human mind are more complex and richer than what discrete reports can capture, and that access to such information is a property of both perception and memory.

Do these results suggest that perception is probabilistic? Clearly, participants have more information in mind than a discrete response. However, this may not be sufficient to conclude that perception is fully probabilistic. Rahnev et al.^[15]^ highlighted the possibility that perception might only be probabilistic in a “weak” sense: People might have a discrete value in mind plus a sense of confidence about their perception. However, one piece of evidence from our findings argues against the weak probabilistic account. As participants place more bets, they do so in an asymmetric fashion, with later bets shifting the mean towards to the true value. This suggests that participants encode and have access to representations more descriptive than would be provided by a confidence interval.

The current findings suggest that perceptual reports have probabilistic properties. A distinct, but equally important question is whether this entails full conscious access to this probabilistic information at a given time. It could be that individual bets (and conscious awareness) are limited to discrete samples that are drawn from an internal probability distribution of information. Evidence of such noisy sampling has been found in many paradigms including decision-making^[28]^, object recognition^[29]^, attention^[30]^, etc. Studies have further argued that access to consciousness is all-or-none^[31,32]^, consistent with Block’s^[1]^ proposal of conscious representations through winner-takes-all processes (but see Karabay et al.^[33]^). However, even if our results were the result of sampling, this means that probabilistic information is perceptually encoded even if there are limits in moment-to-moment accessibility.

Ultimately, whether or not the internal representation can be classified as fully probabilistic or simply probabilistic-like (e.g., a rich collection of samples, etc.), the current paradigm indicates that there is practically useful information not tapped into if only discrete response paradigms are used.

## Data Availability

The datasets generated and analyzed during the current study are available at https://osf.io/vrpbd/

## Footnotes

1 We would like to thank Ned Block for highlighting this possibility and for suggesting a way to test it.

## Notes

https://osf.io/vrpbd/

### Competing Interest Statement

The authors have declared no competing interest.

