## Supplemental Materials for "Perception is Rich and Probabilistic"

### Supplementary Materials

To better understand what underlying mechanisms can produce our data we conducted simulations with explicit predictions on how internal uncertainty is converted into multiple bet responses. This is adapted from what was done previously for our working memory studies[1], but with the parameters updated for the current dataset. We simulated a million trials where on each trial the first response was either a random response (10% of trials) or was drawn from a Gaussian (90% of trials) with a standard deviation sampled from a higher order distribution with a mean precision of 21° and a standard deviation in precision of 7°. These parameters were the best fitting variable precision model[2,3,4] on our data set. We consider four possible means of placing bets based on perceptual / memory representations.

#### Model A: Discrete-estimate stacking

First, we simulate what would be the trends across bets when participants simply stack subsequent bets on bet 1 (with some small added random noise). Bet 1 is the response error generated by the variable precision model. As is evident in Figure S1 (panel a), neither individual errors nor cumulative errors are expected to change across bets under this scenario. This would be true regardless of the parameters used, e.g., changing the guess rate or perceptual *SD* mean or standard deviation changes the magnitude of error across all bets equally. It would not make sense to add a memory component to this model since the initial bet remains on-screen for participants to stack on top of for successive bets.

#### Model B: Confidence-dependent spreading of bets

Model B assumes that participants have and use an estimate of trial-by-trial uncertainty to influence the amount of spread around bet 1. This could occur due to participants having a discrete representation plus some sense of confidence[5]. Here we used the true *SD* for that trial (different trials have different *SDs* thanks to variable precision) provided the basis for spreading bets. This relation was kept linear and at worsening scale. We assigned a Gaussian spread around bet 1 to pick a bet 2. This spread we arbitrarily picked to be a percentage (40%) of the perceptual *SD* of the trial (picking other percentages does not change the trends, only the magnitude of error. To mimic the decay of memory, bet spread thereon will increase by 2° by each bet, i.e., if it was 25° on bet 2, it would be 27° on bet 3, 29° on bet 4, etc. This increase is also somewhat arbitrary, but regardless of how the bet spread is widened per bet, cumulative errors will always increase while individual errors also increase. Even if we removed this memory component and had the bet spread be constant, cumulative error with this model still *never* decreases.

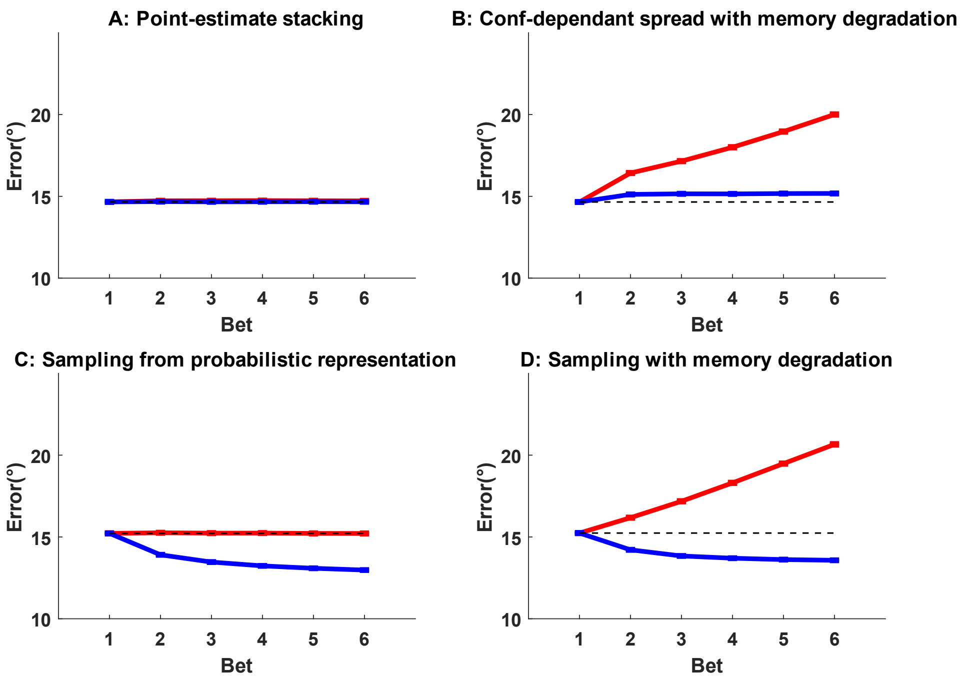


Figure S1. Each panel represents a different mechanism of converting memories into bets. Note that benefits to cumulative error were only seen in the case of sampling.

### Model C: Sampling without degrading information

The previous models have assumed that participants report optimally from the encoded information. However, there are reasons to doubt this assumption. For one, there are multiple lines of evidence that responses may be more akin to random samples from internal probability distributions rather than optimal or representative summaries of this[6,7]. Similarly, with this model we assume that the stored representation is an uncertainty distribution over shape space and that bets are samples from this internal distribution. Due to sampling noise, bet 1 will no longer always be the mode (or highest point) of the internal distribution. Similarly, we assume that bets 2-6 will also be samples. For simplicity, the bets are not influenced by previous responses, although one could imagine a strategy of over- or under-correcting bets based on previous responses. An implication of the sampling model is that bets will have more spread than the internal distribution. To simulate a bet 1 error magnitude that is roughly equivalent to the other models, the assumed internal distribution is less uncertain (consistent with our argument that reports overestimate the uncertainty in memory). For this model we modified the mean of standard deviation of the Gaussian to be 85% used in Models A/B/C (although note that the trends remain the same, even if we use the original internal uncertainty spread). This model predicts a benefit to cumulative error, as seen in our data. Of course, this should not be taken to imply that our data can only be explained by internal sampling, but it highlights that (unlike the other models) sampling can produce the same qualitative pattern as our observed data as far as the cumulative benefit is concerned.

**Model D: Sampling with degrading information**

The previous model assumes that the information quality used to place the last sample is as good as that used to place the first sample. This is unlikely, given that response/perceptual interference[8,9] and time-based decay will result in reduced information quality for later bets. Similar to Model B, we implemented this by increasing error of later bets by 2° of sampling spread per bet. For example, where bets under Model C for a random trial might be from a Gaussian of mode +9° and standard deviation of 16° for all six bets, this will only be true for bet 1 under Model D, with bet 2 being sampled from a Gaussian of the same mode but standard deviation of 18°, etc. This results in increasing errors for later bets (sloped red line in Figure S1), although having only a small impact on the benefit of cumulative error. This model is able to replicate the qualitative pattern of data seen in our experiments. Note that the slope of the increase in individual errors can easily be modified by specifying how spread changes across bets differently. We apply a linear slope as it is the most basic.
